# A collective modulatory basis for multisensory integration in *C. elegans*

**DOI:** 10.1101/424135

**Authors:** Gareth Harris, Taihong Wu, Gaia Linfield, Myung-Kyu Choi, He Liu, Yun Zhang

## Abstract

In the natural environment, animals often encounter multiple sensory cues that are simultaneously present. The nervous system integrates the relevant sensory information to generate behavioral responses that have adaptive values. However, the signal transduction pathways and the molecules that regulate integrated behavioral response to multiple sensory cues are not well defined. Here, we characterize a collective modulatory basis for a behavioral decision in *C. elegans* when the animal is presented with an attractive food source together with a repulsive odorant. We show that distributed neuronal components in the worm nervous system and several neuromodulators orchestrate the decision-making process, suggesting that various states and contexts may modulate the multisensory integration. Among these modulators, we identify a new function of a conserved TGF-β pathway that regulates the integrated decision by inhibiting the signaling from a set of central neurons. Interestingly, we find that a common set of modulators, including the TGF-β pathway, regulate the integrated response to the pairing of different foods and repellents. Together, our results provide insights into the modulatory signals regulating multisensory integration and reveal potential mechanistic basis for the complex pathology underlying defects in multisensory processing shared by common neurological diseases.

**Author Summary:** The present study characterizes the modulation of a behavioral decision in *C. elegans* when the worm is presented with a food lawn that is paired with a repulsive smell. We show that multiple sensory neurons and interneurons play roles in making the decision. We also identify several modulatory molecules that are essential for the integrated decision when the animal faces a choice between the cues of opposing valence. We further show that many of these factors, which often represent different states and contexts, are common for behavioral decisions that integrate sensory information from different types of foods and repellents. Overall, our results reveal a collective molecular and cellular basis for integration of simultaneously present attractive and repulsive cues to fine-tune decision-making.

## Introduction

An environment is often represented by numerous sensory cues. For example, a tainted food source can produce both attractive and repulsive odorants. In order to better survive, an animal often needs to detect and process simultaneously present sensory cues to make a behavioral decision [1–8]. Because integrating multiple sensory cues generates a more accurate evaluation of the environment, it provides important adaptive values. Multisensory integration is widely observed in both the vertebrate and invertebrate animals. Previous studies using behavioral and psychophysical approaches show that humans and other organisms can integrate an array of sensory stimuli to generate decisions in every-day life [9–11]. One common characteristic of multisensory behavioral responses and decision-making processes is their ability to be modulated by various internal states and contexts, including arousal, sleepiness versus wakefulness, the motivational or nutritional state of the organism, or the level of the reward paired with the stimuli. Neurotransmitters, such as dopamine, serotonin, glutamate, and neuropeptides, mediate many of these neurological effects on decision-making [3, 4, 12–14]. Intriguingly, patients of several neurological diseases, including autism spectrum disorder, Parkinson’s disease, bipolar disorder, depression, schizophrenia, and gambling behaviors, share deficits associated with sensory processing or decision-making when encountering multiple sensory stimuli that evoke certain behavioral choices under normal conditions [15–21]. Together, these studies reveal multisensory integration as a common neuronal and behavioral process modulated by multiple contexts across the animal kingdom and highlight the importance of understanding the underlying mechanisms in normal as well as disease states.

Despite the importance of multisensory integration in animal behavior, our understanding of the underlying signaling mechanisms remains preliminary. The nematode *C. elegans* provides an opportunity to address the question. *C. elegans* feeds on bacteria. A bacterial lawn provides various types of sensory information, including olfactory, gustatory, mechanical, and gaseous cues. The small nervous system (302 neurons) of *C. elegans* generates sensorimotor responses to these modalities [22–33] and many of the responses can be shaped by external and internal contexts that modulate neural activities [4, 34–38]. The *C. elegans* genome encodes the homologues of about 50% of the molecules expressed in the mammalian brains [39], which in combination with a well-defined wiring diagram of the nervous system [40] allows characterizing of the molecular and circuit basis for multisensory integration during decision-making.

Here, we show that *C. elegans* integrates the information from an attractive food lawn and a simultaneously present repellent to generate a decision on leaving. We show that the decision to leave the lawn depends on the attractiveness of the lawn and the concentration of the repellent. We identify specific neurons and a collection of modulatory molecules that promote or suppress the food-repellant integration underlying the lawn-leaving decision. We further demonstrate that the battery of modulatory molecules and neurons act as common modulators to regulate integrated decisions on different foods paired with different repellants. These findings identify conserved neuronal signals that modulate multisensory processing during decision-making and reveal a collective modulatory basis for multisensory integration.

## Results

### *C. elegans* integrates multiple sensory cues to generate a behavioral decision

To establish a behavioral assay for multisensory integration in *C. elegans*, we presented a repulsive odorant, 2-nonanone, to the animals on a small lawn of the *E. coli* strain OP50 (Figure 1A and Experimental Procedures) and assessed the decision of the animals to stay on or leave the lawn over time. Because the OP50 lawn serves as a food source for the worm, under the standard condition *C. elegans* stays on the lawn and leaves only at a low frequency [41, 42]. Meanwhile, 2-nonanone strongly repels *C. elegans* at concentrations ranging from 10% to 100%. The olfactory sensory neuron AWB detects and mediates the avoidance of 2-nonanone [43–45]. We first presented a drop of 10% 2-nonanone close to the edge of an OP50 lawn, on which 15 - 25 young adults acclimatized for one hour (Figure 1A). We found that within a few minutes the animals migrated to the side of the bacterial lawn away from 2-nonanone, stayed on the edge of the lawn before dispersing throughout the lawn without leaving (Figure 1B). This result indicates that *C. elegans* is able to detect and avoid 10% 2-nonanone even on the food lawn, but the repulsion is not strong enough to suppress the retention of the worm by the food lawn. In contrast, when we presented a drop of higher concentration of 2-nonanone to the worms in the same configuration, the worms migrated to the side of the lawn, started to leave the food lawn in a few minutes, and continued to migrate to the edge of the plate away from the repellent within the one-hour time window of the assay (Figure 1B, 1C and S9 Movie). The food-leaving behavior was robustly evoked with 100% 2-nonanone (Figure 1B-1E), under which condition a significant number of worms already left the lawn after 2-nonanone was presented to the worms for 15 minutes (Figure 1C). In addition, it took a similar amount of time for the worms to reach the edge of the lawn that was paired with either 10% or 100% 2-nonanone (Figure 1D). These results show that *C. elegans* integrates the attraction of a food lawn with the repulsion of 2-nonanone to generate a behavioral decision and that increasing concentration of 2-nonanone enhances lawn leaving (Figure 1B and 1C). These findings are consistent with the general rule that governs multisensory integration, where increasing the reliability of a sensory cue, such as increasing the concentration of 2-nonanone, strengthens the weight of the cue in integration [46]. To characterize the regulatory mechanisms underlying multisensory integration, we used 100% 2-nonanone as the repellent for the rest of the study unless otherwise described. We quantified the percentage of the worm outside the OP50 lawn 15 minutes after the assay began, because it was an early time point when wild type started to show a robust leaving decision.

**Fig 1.**
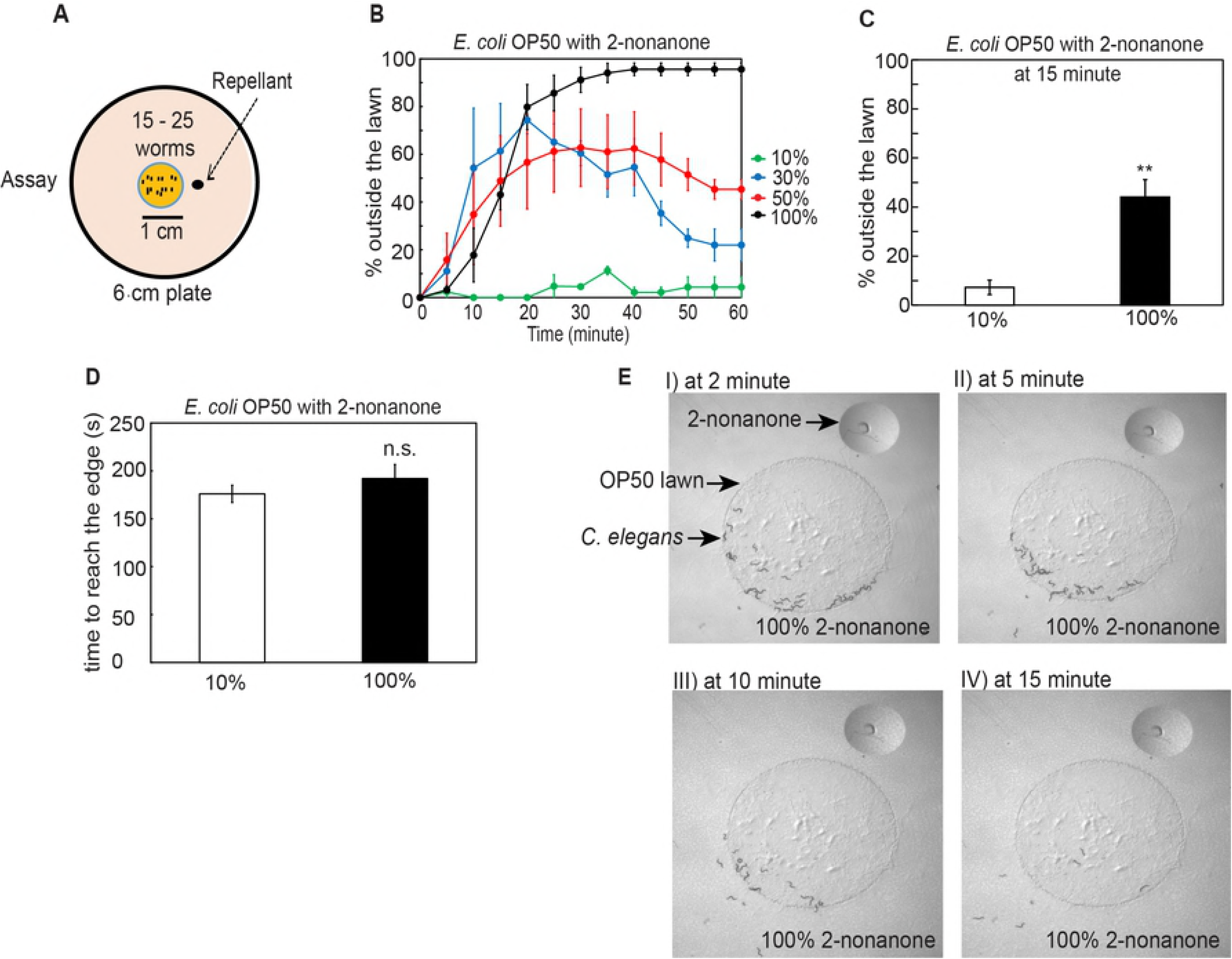
*C. elegans* performs multisensory integration to leave food paired with a repulsive odorant 2-nonanone. A. A schematic of 2-nonanone-dependent food leaving assay.
B. The time course for worms leaving OP50 lawn that is paired with 2-nonanone of different concentrations over 60 minutes, n = 2 assays for 10% and n = 3 assays each for 30%, 50% and 100%.
C. More worms leave the OP50 food lawn paired with 100% 2-nonanone (n = 4 assays) than the OP50 lawn paired with 10% 2-nonanone (n = 5 assays). Bar graph represents the percentage of worms outside the lawn 15 minutes after the assay starts.
D. The time taken for worms to reach the edge of the OP50 food lawn when the lawn is paired with either 10% or 100% 2-nonanone, n = 2 assays each.
E. **I – IV**, Sample images of wild-type animals leaving an OP50 lawn that is paired with 100% 2-nonanone at different time points of the 60-minute assay. For **B-D**, Mean ± SEM, Student’s *t* test, ^⋆⋆^ p ⩽ 0.01, n.s., not significant.

### Sensory neurons that regulate multisensory integration

To characterize how the nervous system regulates the integrated response to the attractive OP50 lawn and the repulsive odorant 2-nonanone, we first probed the amphidal sensory neurons AWB, which mediate avoidance of 2-nonanone via the function of the cGMP-gated channel subunit *tax*-*2* [43]. Exposure to 2-nonanone suppresses the intracellular calcium transients of AWB [44, 45]. Consistently, we found that the transgenic animals that selectively expressed a hyperactive form of an amiloride-sensitive sodium channel MEC-4 that generated necrosis of AWB [43, 47] did not leave the OP50 lawn when 2-nonanone was present (Figure 2A) and that many of the worms remained diffusely distributed on the food lawn by the end of the assay. These transgenic animals were defective in avoiding 2-nonanone in the standard chemotaxis assay (S1 Table and S5 Figure), consistent with previous findings [43]. AWB-killed animals also spent more time to reach the edge of the OP50 lawn when 2-nonanone was present (S2 Table), consistent with the role of AWB in mediating the avoidance of 2-nonanone. Meanwhile, the transgenic animals with genetically killed AWB stayed on OP50 lawn similarly as wild type when 2-nonanone was not present (S3 Table). Together, these results show that AWB regulates the integrated response by mediating the response to the unisensory repellent 2-nonanone.

**Fig 2.**
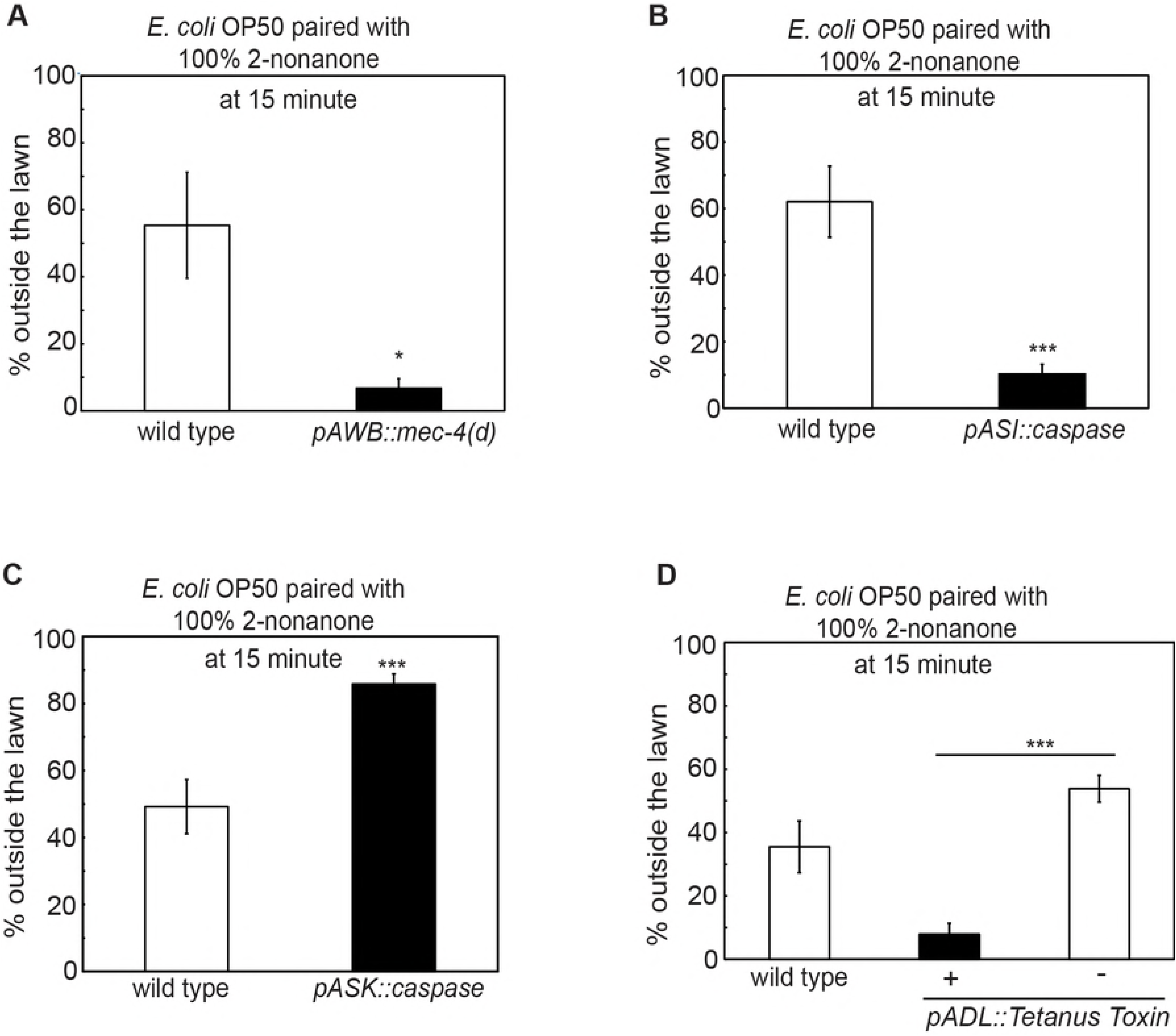
Several sensory neurons modulate 2-nonanone-dependent food leaving. **(A-D)** The transgenic animals that either lack the functional AWB sensory neuron by selectively expressing the gain-of-function isoform of an amiloride-sensitive sodium channel MEC-4 (**A**, *pAWB::mec*-*4(d)*, n = 5 assays each) or lack the ASI sensory neuron by expressing a cell death promoting molecule caspase (**B**, *pASI::caspase*, n = 7 assays for wild type and 8 assays for the transgenic animals) or are defective in the synaptic transmission of the ADL sensory neuron by expressing the tetanus toxin (**D**, *pADL::TeTx*, n = 5 assays for wild type, 4 assays for the transgenic animals, and 3 assays for non-transgenic siblings) display a delayed decision to leave the OP50 lawn paired with 100% 2-nonanone; while the transgenic animals that express caspase in the ASK sensory neuron (**C**, *pASK::caspase*, n = 9 assays each) display a faster decision to leave. Each bar graph reports the average percentage of worms outside the lawn 15 minutes after the assay starts. Mean ± SEM, Student’s -est, ^⋆^ p ⩽ 0.05, ^⋆⋆⋆^p ⩽ 0.001.

Next, we sought additional sensory neurons that regulated the integrated behavioral decision. Previous studies identify several sensory neurons that respond to the smell of the *E. coli* strain OP50 or mediate the behavioral response to the presence or removal of food [22, 44, 48–51]. To examine the potential role of these sensory neurons in our multisensory integration paradigm, we first tested a null mutation *ky4* in *odr*-*7*, which encoded a putative DNA-binding nuclear receptor that specified the function of the AWA sensory neuron [52], a null mutation *p680* in *che*-*1*, which encoded a zinc finger transcription factor required for the development and function of the ASE sensory neuron [53], transgenic animals that selectively expressed a cell-death activator EGL-1 [54] in the AQR, PQR and URX neurons or the CO_2_-sensing BAG sensory neuron [28–30, 55–57]. We also tested transgenic animals selectively expressing a cell-death inducing caspase, or *twk*-*18(gf)* that encoded a constitutively active form of the potassium channel TWK-18 [58], or tetanus toxin that eliminated the synaptic release [59] in the ASI, AWC, ASJ, ADL or ASK neuron [49, 55, 60–64]. We found that all except three of the tested strains were normal. The transgenic animals that contained genetically-killed ASK left the OP50 lawn significantly faster than wild type, and the transgenic animals that contained genetically-killed ASI or expressed the tetanus toxin in ADL left the OP50 lawn significantly more slowly than wild type (Figure 2B-2D and S4 Table). Because the transgenic animals defective in the function of ASK or ASI or ADL are not deficient in avoiding 2-nonanone in our standard chemotaxis assay, in their ability to remain on OP50 lawn when 2-nonanone is not present, as well as in moving to the edge of the OP50 lawn with the presence of 2-nonanone (S1-3 Tables), these results together indicate that the sensory neurons ASK, ASI and ADL modulate how rapidly the behavioral decision to leave the repellent-paired food lawn is made.

### Multisensory integration requires peptidergic and the TGF-β pathways

To characterize the mechanisms underlying multisensory integration of food and 2-nonanone, we examined mutants that were defective in biosynthesis of neurotransmitters. We tested effects of mutating *tph*-*1* that encoded tryptophan hydroxylase required for the production of serotonin [65], *cat*-*2* that encoded tyrosine hydroxylase needed for the synthesis of dopamine [66], *tdc*-*1* that encoded tyrosine decarboxylase required for the synthesis of tyramine and octopamine, or *tbh*-*1* that encoded tyramine beta hydroxylase required for the production of octopamine [67]. Interestingly, all of these mutants exhibited wild-type behavioral decision when they were exposed to 2-nonanone on an OP50 food lawn (Figure 3A-D and S6 Figure). These results show that serotonin, dopamine, tyramine or octopamine are not required for 2-nonanone-dependent food leaving, although these neurotransmitters regulate many food-dependent sensorimotor responses ([32] and references therein).

**Fig 3.**
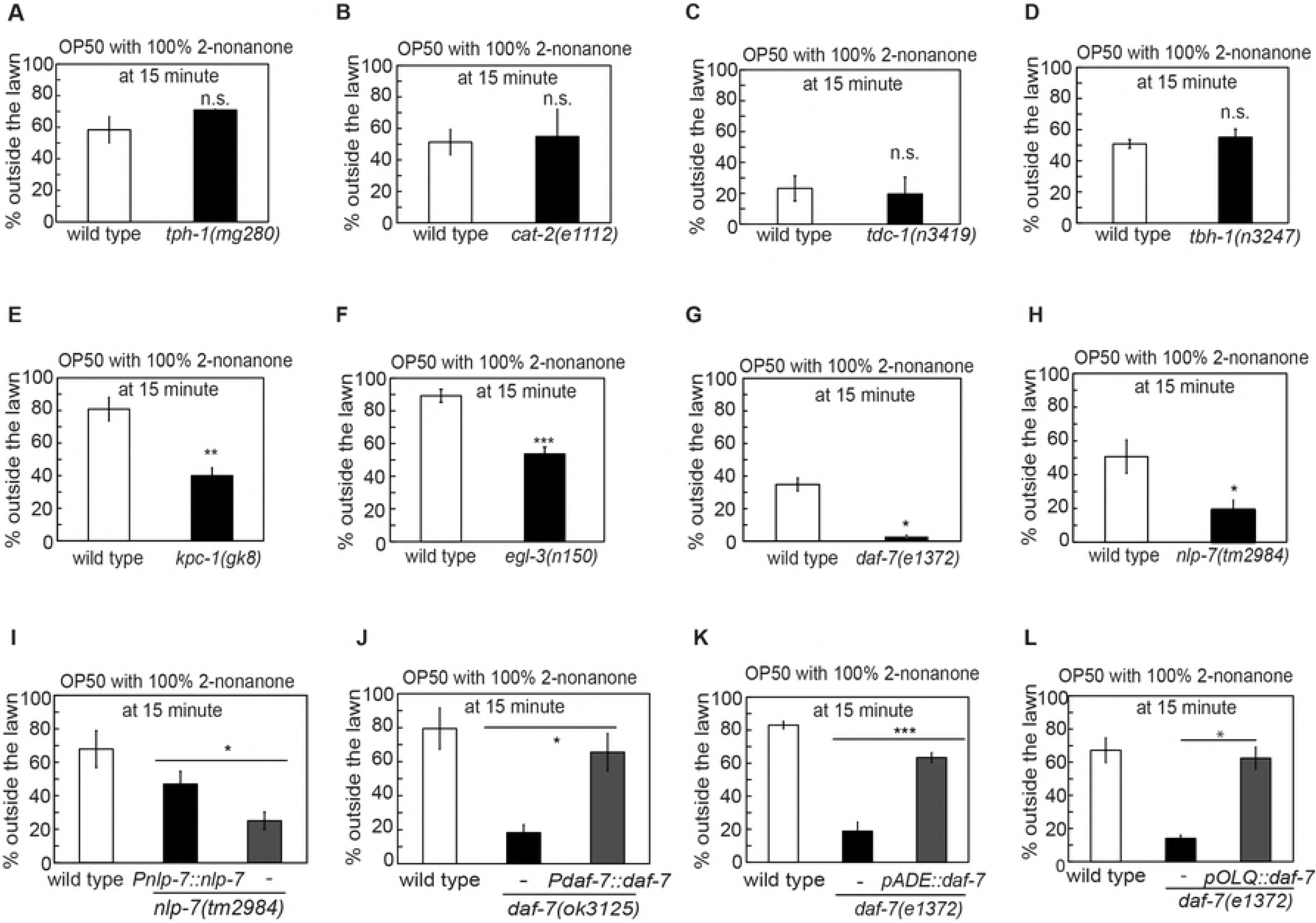
NLP-7 and TGF-β/DAF-7 modulate the decision to leave the OP50 food lawn paired with 2-nonanone. **(A-D)** The mutant animals that are defective in the biosynthesis of the neurotransmitter serotonin (**A**, *tph*-*1(mg280)*, n = 2 assays each), or dopamine (**B**, *cat*-*2(e1112)*, n = 4 assays each), or tyramine and octopamine (**C**, *tdc*-*1(n3419)*, n = 2 assays each), or octopamine (**D**, *tbh*-*1(n3247)*, n = 3 and 4 assays for wild type and *tbh*-*1* mutants, respectively) display a normal decision to leave the OP50 food lawn that is paired with 2-nonanone. **(E-K)** Mutations in the genes encoding the neuropeptide processing enzymes, *kpc*-*1* (**E**, n = 4 and 5 assays for wild type and *kpc*-*1* mutant, respectively), or *egl*-*3* (**F**, n = 4 and 6 assays for wild type and *egl*-*3* mutant, respectively), or a TGF-β -encoding gene *daf*-*7* (**G**, n = 6 and 5 assays for wild type and *daf*-*7* mutant, respectively), or a neuropeptide-encoding gene *nlp*-*7* (**H**, n = 7 and 8 assays for wild type and *nlp*-*7* mutant, respectively) generate a delayed decision to leave the OP50 food lawn paired with 2-nonanone, and expressing the genomic DNA of *nlp*-*7* (**I**, n = 6, 7 and 4 assays for wild type, transgenic animals and non-transgenic siblings, respectively) or *daf*-*7* (**J**, n = 4 assays each for wild type, transgenic animals and non-transgenic siblings) rescues the delayed food leaving phenotype of the respective mutant animals. **K)** Expressing the wild-type *daf*-*7* cDNA in the sensory neurons ADE rescues the delayed decision in the *daf*-*7(e1372)* mutant animals, n = 4 assays each for wild type, transgenic animals and non-transgenic siblings, respectively. **(L)** Expressing the wild-type *daf*-*7* cDNA in the sensory neurons OLQ also rescues the delayed decision in the *daf*-*7(e1372)* mutant animals, n = 3 assays for wild type, 3 assays for transgenic animals and 2 assays for non-transgenic siblings, respectively. Each bar graph reports the average percentage of worms outside the lawn 15 minutes after the start of the assay, mutants are compared with wild-type animals and transgenic animals are compared with non-transgenic siblings using Student’s *t*-*test*, ^⋆^ p ≤ 0.05, ^⋆⋆^ p ≤ 0.01, ^⋆⋆⋆^p ≤ 0.001, n.s., not significant.

Next, we examined the function of neuropeptide-encoding genes. We first found that mutations in the *kpc*-*1(gk8)* and *egl*-*3(n150)* that disabled two of the four known peptide pre-processing enzymes in *C. elegans* [68–70], delayed the decision to leave the food lawn that was paired with 2-nonanone (Figure 3E and 3F), suggesting the modulatory role of peptides or growth factors in promoting the integrated decision to leave the tainted food lawn. Next, we screened many mutations in genes encoding peptides or growth factors. We focused on the available mutations that did not generate any gross defect in either development or locomotion and identified three mutations that significantly altered the wild-type decision. The canonical mutations, *e1372*, or a deletion, *ok3125*, in *daf*-*7* that encoded a TGF-β ligand that regulated development, metabolism and host-pathogen recognition [71–73], significantly delayed the decision to leave the OP50 lawn tainted with 2-nonanone (Figure 3G and 3I). A deletion mutation *tm2984* in *nlp*-*7*, which encoded a neuropeptide-like protein that regulated stress response, egg-laying, life span and modulation of aversive responses to noxious stimuli [74–77], similarly delayed the decision to leave the lawn (Figure 3H). However, the mutations in *daf*-*7 or nlp*-*7* did not generate any detectable defect in the chemotactic response to 2-nonanone alone, or the tendency to leave the OP50 lawn when no repellent was present, or the ability to move to the edge of the lawn when 2-nonanone was present (S1-3 Tables). In addition, expressing the genomic fragment containing the regulatory and coding regions of *daf*-*7* or *nlp*-*7* fully rescued the defect of the respective mutant animals in making the decision to leave the lawn that was paired with 2-nonanone (Figure 3I and 3J). Together, these results indicate that TGF-βDAF-7 and NLP-7 promote the food-leaving decision when 2-nonanone is present.

### A new function of the TGF-βDAF-7 canonical pathway in multisensory integration

The *C. elegans* TGF-β/DAF-7 regulates several physiological processes through the conserved type I and type II TGF-β receptor, DAF-1 and DAF-4, respectively [78, 79]. DAF-7 is found in the sensory neurons OLQ, ADE and ASI, all of which are implicated in sensing bacteria [23, 71, 72, 80, 81]. DAF-7 produced by ASI regulates the satiety-induced quiescence, the entry into an alternative developmental stage under the environmental stress, and the modulation of the lifespan by dietary restriction, and responds to the abundance of food [72, 80–83]. The expression of *daf*-*7* is induced in the sensory neuron ASJ upon exposure to pathogenic bacteria and DAF-7 in ASJ regulates the avoidance of the pathogen through DAF-1 and DAF-4 receptors [71]. In addition, through DAF-1 and DAF-4, DAF-7 regulates metabolism and fat accumulation [73]. Here, we showed that mutating *daf*-*7* delayed the decision to leave the food lawn when 2-nonanone was present (Figure 3G). To identify the source of the DAF-7 signal regulating multisensory integration, we tested the transgenic animals that selectively expressed a wild-type *daf*-*7* cDNA in subsets of *daf*-*7*-expressing neurons in *daf*-*7* mutant animals for potential rescuing effects. We found that expressing *daf*-*7* selectively in ASI (*Pstr*-*3::daf*-*7* [71]) did not rescue the defects in the integrated response; but expressing *daf*-*7* in ADE (*Pcat*-*2::daf*-*7*) or OLQ (*Pocr*-*4::daf*-*7*) sensory neurons using cell-selective promoters [84–86] rescued the delayed leaving phenotype in the *daf*-*7(e1372)* mutant animals (Figure 3K and 3L, S7 Figure). In addition, we found that the canonical mutation in the type I and type II TGF-β receptor, *daf*-*1(m40)*, similarly delayed the decision to leave the OP50 lawn paired with 2-nonanone (Figure 4A). Expressing either the genomic DNA fragment of *daf*-*1* or the *daf*-*1* cDNA selectively in the interneurons RIM and RIC (*Pdaf*-*1::daf*-*1* or *Ptdc*-*1::daf*-*1;* [71, 73]) fully rescued the defect in the *daf*-*1(m40)* mutant animals (Figure 4B and 4C), while expressing *daf*-*1* in sensory neurons (*Pbbs*-*1::daf*-*1* or *Posm*-*6::daf*-*1*; [71]) was not sufficient to rescue (Figure 4D and S8 Figure). Together, these results indicate that the TGF-β/DAF-7 signal produced by the ADE or the OLQ sensory neurons acts through the type I TGF-β receptor DAF-1 in RIM and/or RIC neurons to promote repellent-dependent leaving of the food lawn.

**Fig 4.**
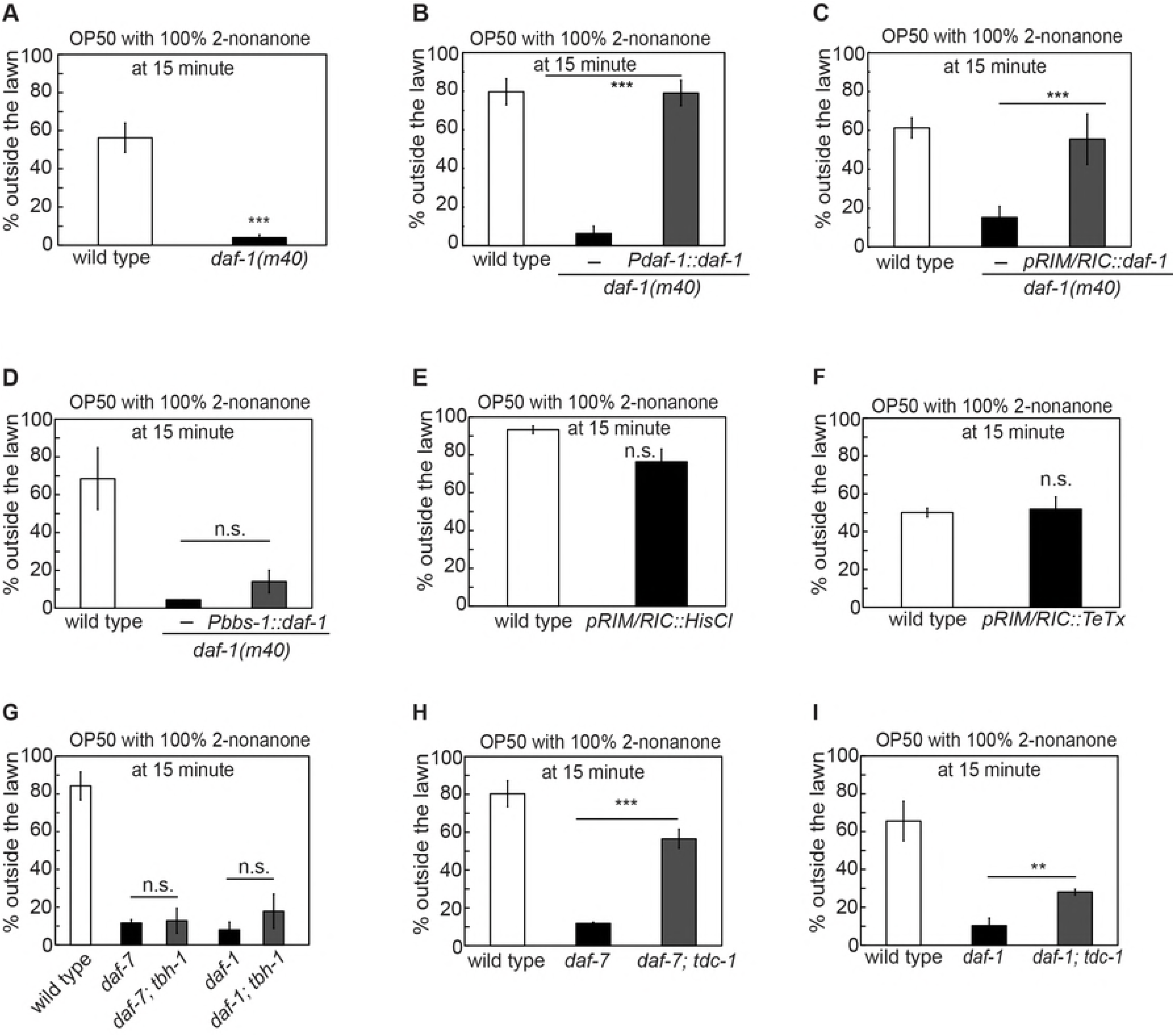
The TGF-β receptor DAF-1 acts in the RIM and RIC neurons to mediate 2-nonanone-dependent food leaving. **(A-D)** Mutating *daf*-*1* that encodes the type I TGF-β receptor delays the decision to leave the OP50 lawn paired with 100% 2-nonanone (**A**, n = 8 assays each), and expressing the genomic DNA of *daf*-*1* (**B**, n = 6 assays each) or a wild-type *daf*-*1* cDNA in the RIM and RIC neurons (**C**, n = 6, 6 and 5 assays for wild type, transgenic animals and non-transgenic siblings, respectively) in the *daf*-*1(m40)* mutant animals rescues the delayed decision, but expressing wild-type *daf*-*1* in the sensory neurons (**D**, n = 3, 5 and 2 assays for wild type, transgenic animals and non-transgenic siblings, respectively) does not rescue. Mutants are compared with wild type and transgenic animals are compared with non-transgenic siblings with Student *t*-test. **(E-F)** Inhibiting the activity of the RIM and RIC neurons by selectively expressing a histamine-gated chloride channel (**E**, n = 2 and 4 assays for wild type and transgenic animals, respectively) or blocking the synaptic release from these neurons by selectively expressing the tetanus toxin (**F**, n = 2 assays each) does not alter the decision to leave the OP50 food lawn paired with 100% 2-nonanone. Transgenic animals are compared with wild type. **(G-I)** Removing octopamine signaling in the *daf*-*7(e1372)* or *daf*-*1(m40)* mutants with a mutation that disrupt biosynthesis of octopamine *tbh*-*1(ok1196)* does not suppress the delayed leaving from the OP50 lawn paired with 100% 2-nonanone (**G**, n = 6 assays for wild type; n = 4 assays for *daf*-*7* mutants; n = 2 assays for *daf*-*1* mutants; n = 3 assays for *daf*-*7;tbh*-*1* double mutants; n = 2 assays for *daf*-*1*;*tbh*-*1* double mutants), but removing the tyramine and the octopamine signals with the mutation in *tdc*-*1(ok914)* in either the *daf*-*7(e1372)* (**H**, n = 6, 5 and 4 assays for wild type, *daf*-*7* mutants and *daf*-*7;tdc*-*1* double mutants, respectively) or the *daf*-*1(m40)* (**I**, n = 5, 4 and 5 assays for wild type, *daf*-*1* mutants and *daf*-*1;tdc*-*1* double mutants, respectively) mutant animals suppresses the delay-decision phenotype in either of the mutant animals. Double mutants were compared with the respective single mutants using student’s *t* test. Each bar graph reports the average percentage of worms outside the lawn 15 minutes after the start of the assay. Mean ± SEM, ^⋆⋆^ p ⩽ 0.01, ^⋆⋆⋆^ p ⩽ 0.001, n.s.; not significant.

To further interrogate the role of the RIM/RIC neurons in multisensory integration, we examined the transgenic animals that expressed a histamine-gated chloride channel in the RIM and RIC neurons under the histamine-treated condition [87] or the transgenic animals that expressed tetanus toxin [59] in RIM and RIC. We found that these transgenic animals were normal in leaving the OP50 lawn when 2-nonanone was present (Figure 4E and 4F). Since neither the *tdc*-*1(n3419)* mutant animals that lacked tyramine and octopamine nor the *tbh*-*1(n3247)* mutant animals that lacked octopamine is defective in their decisions to leave the OP50 lawn paired with 2-nanone (Figure 3), together, our results suggest that RIM/RIC and the release of the neurotransmitter tyramine and octopamine from these neurons may be suppressed during the integrated response to the simultaneously present food lawn and 2-nonanone. To further interrogate the role of tyramine or octopamine signaling in the *daf*-*7*- and *daf*-*1* -dependent integrated response, we tested how removing tyramine and/or octopamine affects the delayed food leaving in the *daf*-*7(e1372) or daf*-*1(m40)* mutant animals. Interestingly, both of the *daf*-*1(m40); tbh*-*1(ok1196)* and the *daf*-*7(e1372); tbh*-*1(ok1196)* double mutant animals [19] behaved like the *daf*-*1(m40)* and the *daf*-*7(e1372)* single mutants, respectively (Figure 4G). In contrast, the mutation in *tdc*-*1(ok914)* strongly suppressed the delayed decision phenotype in both *daf*-*7(e1372)* and *daf*-*1(m40)* mutant animals (Figure 4H and 4I). While TDC-1 is needed for the production of tyramine and octopamine in both RIM and RIC, TBH-1 is only needed for the biosynthesis of octopamine in RIC [67]. Together, these results show that the TGF-β/DAF-7 regulates the decision between staying on a food lawn versus avoiding a repellent through the canonical signaling pathway and that the DAF-7 peptidergic signal produced from ADE or OLQ inhibits the tyramine neurotransmission of RIM and/or RIC to promote the decision to leave the food-lawn that is paired with 2-nonanone.

### Different interneurons play opposite roles in multisensory integration

To better characterize the neural circuits underlying multisensory integration, we probed the potential interneurons that regulated the decision between staying on the food lawn versus avoiding 2-nonanone. We focused on the interneurons AIY, AIB, and the command interneurons, all of which regulate locomotion [22, 88]. AIY and AIB are also the major interneurons postsynaptic to the sensory neurons that respond to the bacterial food or the repellent 2-nonanone [40]. To disrupt the function of AIY, we selectively expressed in AIY a gain-of-function isoform of a potassium channel TWK-18 [58] to inhibit the activity of AIY (*Pttx*-*3::twk*-*18(gf)*) or the tetanus toxin (*Pttx*-*3::TeTx*) to block synaptic release. We also tested the *ttx*-*3(mg158)* mutants that failed to develop AIY interneurons [89]. All three mutations delayed the decision to leave the lawn (Figure 5A - 5C). However, these manipulations do not disrupt the ability to reach the edge of the food lawn during 2-nonanone-dependent food leaving, to avoid 2-nonanone alone, or to stay on OP50 lawn when no repellent was present (S1-3 Tables). In contrast, selectively expressing the tetanus toxin in the AIB interneurons or treating the transgenic animals expressing the histamine-gated chloride channel in AIB with histamine did not significantly alter the decision to leave the OP50 lawn that was paired with 2-nanonone (Figure 5D and 5E). Together, these results indicate that the activity and the synaptic output of the AIY interneurons promote the decision to leave the food lawn paired with 2-nonanone, while AIB is dispensable for the decision-making. Next, we examined transgenic animals that expressed the tetanus toxin with the *nmr*-*1* promoter or the *glr*-*1* promoter. The *nmr*-*1* promoter is expressed in a few command interneurons including AVA, AVB, AVD, AVE and PVC, while the *glr*-*1* promoter is expressed in several head motor neurons in addition to the *nmr*-*1*-expressing interneurons [90]. Interestingly, both transgenic lines left the 2-nonanone paired food lawn more than wild type (Figure 5F and 5G). However, these transgenic animals are normal in 2-nonanone avoidance in the absence of food or spontaneous food leaving. They also do not reach the edge of the lawn more rapidly than wild type (S1-3 Tables). Together, these results show that different downstream neurons modulate the decision to leave the repellent tainted food lawn in opposite ways by promoting or inhibiting the decision-making process. These neurons may act as the convergent sites to process multiple sensory signals in order to generate specific behavioral outputs.

**Fig 5.**
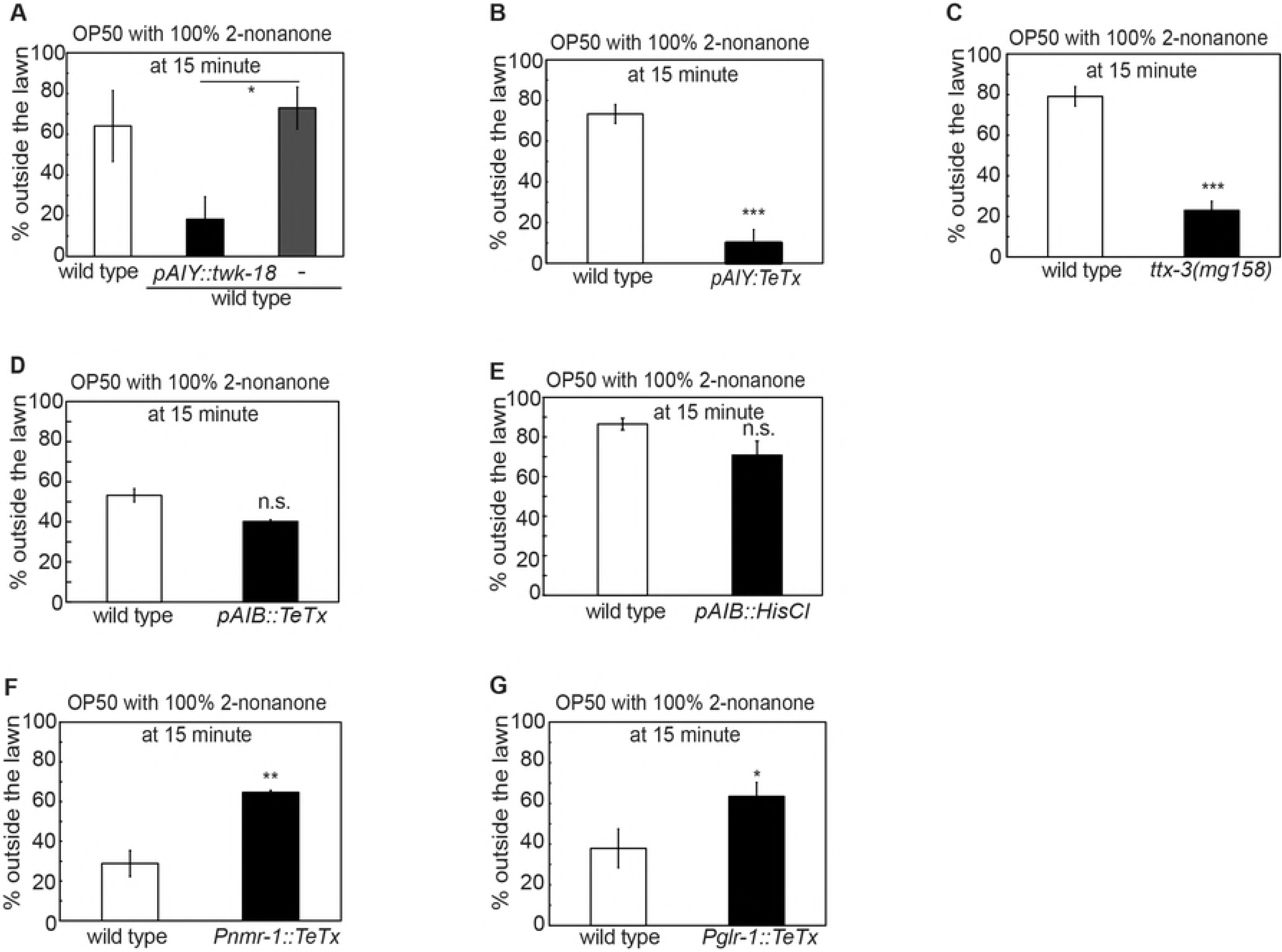
Downstream circuit that regulates 2-nonanone-dependent food leaving. **(A-C)** Inhibiting the activity of the AIY interneuron by expressing the gain-of-function isoform of the potassium channel TWK-18 (**A**, *Pttx*-*3::twk*-*18(gf)*, n = 3 assays each) or by blocking the synaptic outputs of AIY by expressing tetanus toxin (**B**, *Pttx*-*3::TeTx*, n = 4 assays each), or the mutation *ttx*-*3(mg158)* that generates development defects in AIY (**C**, n = 3 assays each) delays the decision to leave the OP50 lawn paired with 100% 2-nonanone. **(D, E)** Selectively expressing tetanus toxin (**D**, *Pinx*-*1::TeTx*, n = 2 assays each) or the inhibitory HisCl channel (**E**, *Pinx*-*1::HisCl*, n = 3 and 4 assays for wild type and transgenic animals, respectively) in the AIB interneuron does not significantly alter the lawn-leaving decision, when the OP50 lawn is paired with 100% 2-nonanone. **(F, G)** Blocking synaptic outputs from the *nmr*-*1*-expressing neurons (**F**, *Pnmr*-*1::TeTx*, n = 3 assays each) or the *glr*-*1*-expressing neurons (**G**, *Pglr*-*1::TeTx*, n = 4 and 3 assays for wild type and transgenic animals, respectively) enhanced the 2-nonanone-dependent lawn leaving. Each bar graph reports the average percentage of worms outside the lawn 15 minutes after the start of the assay. Mean ± SEM, mutants are compared with wild-type animals with student’s *t* test, transgenic animals are compared with non-transgenic siblings with student’s t-test, ^⋆^ p ⩽ 0.05, ^⋆⋆^ p ⩽ 0.01, ^⋆⋆⋆^ p ⩽ 0.001, n.s., not significant.

### Multisensory integration is regulated by a common set of modulators

Next, we asked whether the molecular and circuit mechanisms underlying the integrated response to the OP50 food lawn paired with 2-nonanone were shared by the integrated responses to different pairing of attractive foods and repulsive odorants. We paired the OP50 lawn with various repellants, including 100% 1-octanol and 100% benzaldehyde. While benzaldehyde is attractive at low concentrations [24], 100% benzaldehyde strongly repels *C. elegans* in a way that is dependent on the function of the sensory neuron AWB [91–93]. We found that a drop of 100% benzaldehyde first repelled the animals to the edge of the OP50 food lawn and then in about 10-15 minutes started to repel the animals off the food lawn (Figure 6A and S10 Movie). Interestingly, 1-octanol failed to stimulate food leaving under our experimental conditions (Figure 6A). We also paired a lawn of *Comamonas sp* with 100% 2-nonanone. *Comamonas* is an attractive food source for *C. elegans* [41]. We found that pairing a *Comamonas* bacterial lawn with 100% 2-nonanone repelled *C. elegans* off the lawn similarly as the OP50 lawn paired with 2-nonanone (Figure 7A). Interestingly, we found that several modulators, particularly TGF-β/DAF-7, the TGF-β receptor DAF-1, and the sensory neurons ASK, that regulated the integrated response to an OP50 lawn paired with 100% 2-nonanone also similarly regulated the integrated response to OP50 lawn paired with 100% benzaldehyde and the integrated response to the *Comamonas* lawn paired with 100% 2-nonanone (Figure 6 and 7). Together, these results indicate that a common set of modulators and signaling mechanisms regulates integrated behavioral decision on whether to leave or stay on an attractive food lawn paired with an odorant repellent.

**Fig 6.**
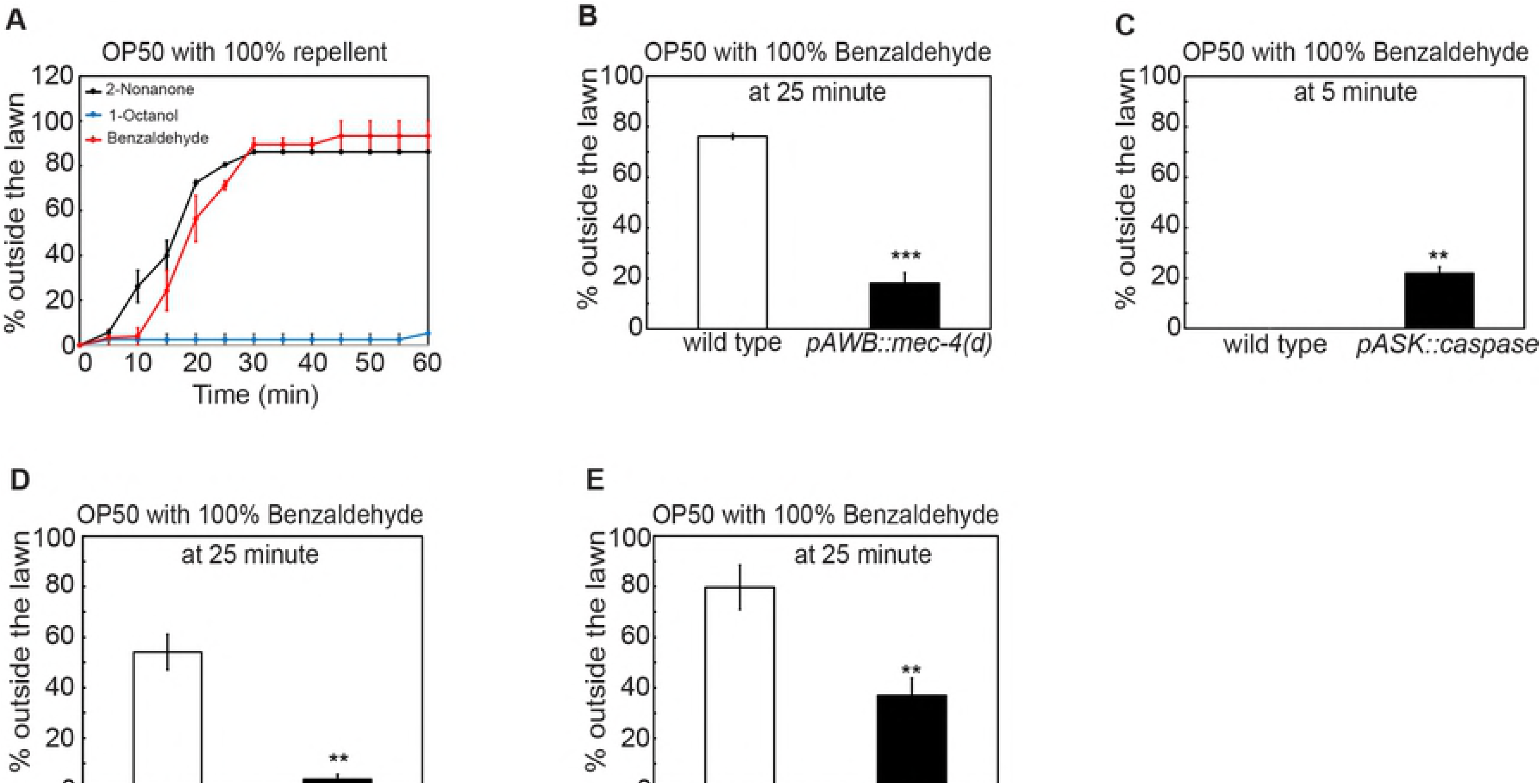
Integrated response to a repellent-paired food lawn is regulated by a common set of factors. **(A)** Wild-type animals also leave the lawn of OP50 paired with 100% benzaldehyde; in contrast, paring an OP50 lawn with either 100% octanol does not repel worms (n = 2 assays for each condition). **(B-E)** Genetic ablation of the sensory neuron AWB (**B**, *pAWB::mec*-*4(d)*, n = 3 assays each) or ASK (**C**, *pASK::caspase*, n = 3 assays each) or mutating the genetic components of the TGF-βDAF-7 pathway (**D**, *daf*-*7(e1372)*, n = 3 assays each; **E**, *daf*-*1(m40)*, n = 4 assays each) alters the decision to leave the benzaldehyde-paired OP50 lawn. Each bar graph reports the average percentage of worms outside the lawn 25 minutes (**B, D, E)** or 5 minutes (**C**) after the start of the assay. Mean ± SEM, mutants are compared with wild-type animals with Student’s *t* test, ^⋆⋆^ p ⩽ 0.01, ^⋆⋆⋆^ p ⩽ 0.001, n.s., not significant.

**Fig 7.**
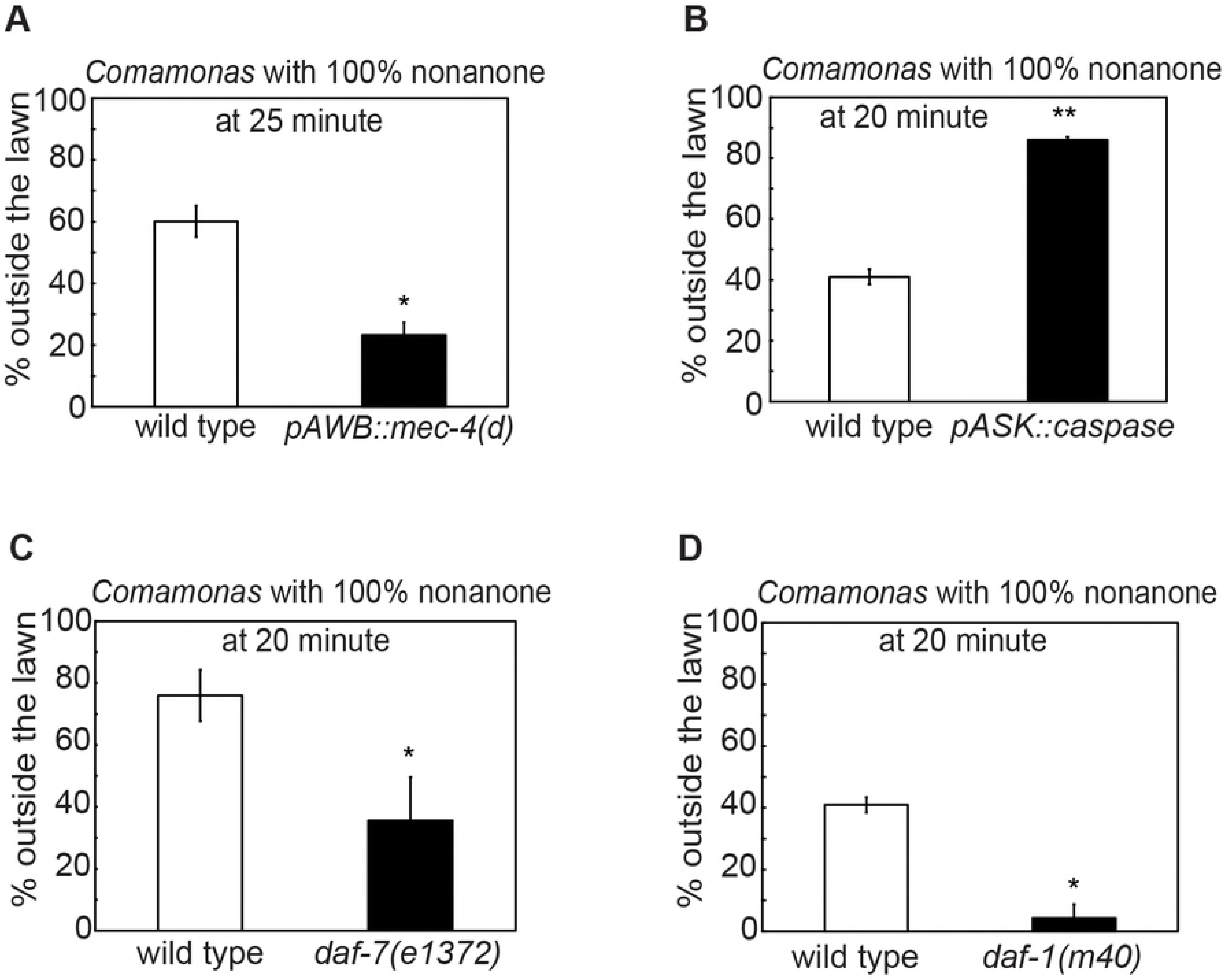
Integrated response to a repellant-paired food lawn requires a common set of factors. **(A-D)** Genetic ablation of the sensory neuron AWB (**A**, *pAWB::mec*-*4(d)*, n = 2 assays each) or ASK (**B**, *pASK::caspase*, n = 2 assays each), or mutating the genetic components of the TGF-βDAF-7 pathway (**C**, *daf*-*7(e1372)*, n = 4 assays each; **D**, *daf*-*1(m40)*, n = 2 assays each) alters the decision to leave the 2-nonanone-paired *Comamonas* lawn. Each bar graph reports the average percentage of worms outside the lawn 25 minutes (**A**) or 20 minutes (**B-D**) after the start of the assay. Mean ± SEM, mutants or transgenic animals are compared with wild type with Student’s *t* test, ^⋆^ p ⩽ 0.05, ^⋆⋆^ p ⩽ 0.01.

## Discussion

Many organisms can combine information from multiple simultaneously present sensory cues to regulate behavioral outputs [1–8, 32]. While the importance of integrated behavioral responses to multiple sensory stimuli is appreciated, the underlying molecular and signaling mechanisms are not well understood. Using our behavioral paradigm for multisensory integration, we characterize the modulators and signaling pathways that regulate a decision to leave a food lawn that is paired with a repulsive odorant. These findings reveal a new function of a conserved TGF-β that modulates decision-making by regulating the tyramine signal from a set of central neurons. Our results elucidate a set of common molecular and neuronal factors that mediate decision-making when the worm is presented with different pairs of stimuli generated by an attractive food and a repulsive odorant (Figure 8).

### Specific sensory neurons regulate multisensory integration

One potential mechanism to regulate a coherent behavioral response to multiple simultaneously present sensory cues is to utilize sensory neurons that are capable of perceiving some or all of the cues. These types of sensory responses can involve either activation or inhibition of certain sensory neurons that detect distinct stimuli. Worms are capable of sensing both food signals and a range of repulsive cues ([32] and the references therein). Here, we characterize the functions of several sensory neurons in regulating the repellent-dependent food leaving when *C. elegans* is exposed to an attractive food lawn concurrently with a repulsive odorant. We confirm the requirement of the AWB sensory neuron that is known to sense repellents, including 2-nonanone and 100% benzaldehyde [43, 45, 91]. AWB also responds to bacterial food [44, 51]. Previous studies identify the role of AWB in promoting food leaving under malnourished conditions [94], suggesting the involvement of AWB in integrating the nutritional state with the food signals. Thus, AWB may regulate the integrated response by simultaneously processing food smells and repulsive odorants.

Interestingly, we also uncover a novel role of three sensory neurons that modulate the decision to leave a food paired with a repulsive odorant. We show that the ASK sensory neuron suppresses the integrated food-leaving decision, while the ASI and ADL sensory neurons promote it. In contrast, we find that several other sensory neurons previously implicated in mediating responses to food-related cues, including AWA, ASE, AWC, ASJ, BAG, AQR, PQR or URX [44, 48, 49, 51, 71], are dispensable in regulating the integrated behavioral response to the food lawn paired with 2-nonanone (S4 Table). In addition, our results suggest that the effect of removing either ASK or ASI or ADL on the decision-making does not result from the altered chemotactic response to 2-nonanone as a unisensory cue or to the OP50 bacterial lawn alone (S1-3 Tables). These results together reveal a specific function of ASK, ASI and ADL in integrating the food signals with the repellent to generate a decision between two sensory cues of opposing valence.

Previous studies show that ASK and ASI sensory neurons respond to *E. coli* OP50 by changing the intracellular calcium levels [51, 95]. Both ASI and ASK are involved in evaluating the food environment. ASI also mediates the balance between food intake and fat storage, as well as experience-dependent changes in food response [37, 73, 80–82, 96]. ASK regulates responses to pheromones and plays a role in food leaving in mutants that are significantly food-deprived and modulates locomotion during pre-exposure to unpredictable food environments [49, 63, 64, 94]. The ADL sensory neuron has been shown to modulate the responses to octanol, pheromone and the preference for certain food odors [64, 77], suggesting that ADL mediates various context-dependent sensory responses to modulate behavior and decision-making. In our present study, we propose that ASI, ADL and ASK neurons represent the strength of the food signals in an antagonistic manner to mediate a balanced behavioral decision between an attractive food and a repulsive odorant.

### Neuropeptides and growth factors modulate multisensory integration

Although previous studies characterize the function of neuromodulators, including neuropeptides and growth factors, in modulating olfactory responses and nutrition-dependent state of the nervous system, how neuromodulatory signals regulate a behavioral decision that integrates cues of opposing values is not well understood. Here, we characterize the role of neuromodulatory molecules, including a conserved TGF-β, in modulating the decision to leave a food lawn when a repulsive odorant is presented together with the lawn.

Neuropeptides and growth factors have been implicated in the context-dependent modulation of several sensorimotor responses in *C. elegans* [3, 34, 77, 96–98]. Here, we identify the neuropeptidergic signaling mechanisms based on the examination of the neuropeptide processing mutants, *egl*-*3* and *kpc*-*1*, and identification of the NLP-7 peptide and the TGF-β/DAF-7 that modulate the decision to leave a food lawn paired with a repulsive odorant. Mutating *nlp*-*7* or *daf*-*7* delayed the decision to leave the lawn. In contrast, mutating genes that encode several other peptides that are expressed in different sensory neurons and have been shown to signal contextual cues or previous experience or food signals, including *ins*-*6*, *ins*-*7*, *nlp*-*1*, *nlp*-*9*, *nlp*-*24* and *flp*-*19* [62, 77, 96, 97, 99], does not have a significant effect (S4 Table), suggesting a specific function of *nlp*-*7* and *daf*-*7* in regulating the decision-making process. *nlp-7* is expressed in several amphidial sensory neurons that respond to contextual cues and NLP-7 delays the acute avoidance of a noxious stimulus, 1-octanol [74–77, 100]. This effect is in contrast with that of mutating *nlp*-*7* in the integrated behavioral response, where NLP-7 promotes the decision to leave the food in order to avoid the repulsive odorant. Our results together with the previous findings characterize distinct functions of the NLP-7 neuropeptide in regulating multisensory integration versus context-dependent avoidance of noxious stimuli.

Previous studies show that the DAF-7 pathway regulates dauer formation, food intake, fat storage, as well as avoidance of pathogenic bacteria after prolonged exposure. The functions of DAF-7 in these physiological events depend on its expression in the sensory neurons ASI and/or ASJ [71–73]. Here, we show that DAF-7 promotes the decision to leave the food lawn paired with a repulsive odorant and that different from its previously identified role, the function of DAF-7 in regulating multisensory decision depends on the expression of *daf*-*7* in either the ADE or the OLQ sensory neurons. Our results are the first to characterize the function of *daf*-*7* produced by ADE or OLQ. ADE is one of the dopaminergic neurons in the worm nervous system [101]. However, we did not see any phenotype in the *cat*-*2* mutants that were defective in dopamine synthesis (Figure 3), suggesting that the function of ADE in regulating the integrated response to food and repellent is independent of dopamine. Both ADE and OLQ have been previously implicated in mechanosensation – OLQ is implicated in sensing the gentle touch delivered to the nose and ADE contributes to the slowing response when a worm enters a bacterial lawn [23, 31, 102]. We propose that ADE and OLQ regulate the integrated response to a food lawn paired with a repellent by representing the mechanical stimulus that a worm senses from the food lawn.

We further show that the canonical TGF-β receptor DAF-1 acts in the interneurons RIM and RIC to regulate the decision to leave the food lawn paired with a repulsive odorant. Interestingly, inhibiting the activity of RIM and RIC, or blocking the synaptic outputs of these neurons, or disrupting the biosynthesis of the common neurotransmitter of these neurons, tyramine, does not significantly change the decision-making process. However, disrupting the production of tyramine, but not octopamine, in these neurons suppresses the slow-decision phenotype in the *daf*-*1* or *daf*-*7* mutant animals (Figure 3 and 4). Together, these results indicate that the DAF-7/DAF-1 pathway promotes the decision to leave by inhibiting the tyramine signaling from these interneurons. This regulatory mechanism of DAF-7 is reminiscent of that in feeding, where DAF-7 promotes the pumping rate by inhibiting the output from the RIM and/or RIC neurons. However, different from the function of DAF-7 in regulating feeding that is dependent on tyramine and/or octopamine [73], DAF-7 modulates the signal of tyramine, but not octopamine, to regulate the decision to leave a food paired with the repellant 2-nanonone.

The RIM and RIC neurons have been previously implicated in various sensorimotor responses, as well as the context-dependent locomotory and feeding behaviors [67, 73, 87, 100, 103]. Our results that characterize RIM/RIC as the downstream neurons of the TGF-β/DAF-7 signal in regulating the decision to leave a tainted food lawn further reveal RIM/RIC as one of the central sites where different sensory signals converge to generate appropriate behavioral outputs. Previously, TGF-β signals have been implicated in various neuronal functions, including learning and memory, neural plasticity in the forms of LTP and LTD, synaptic formation, dendritic development, and regulation of the function of the neural-muscular junctions [104–107]. Defects in TGF-β pathways have been implicated in the pathology of neurological disorders, such as schizophrenia, depression, anxiety and Alzheimer’s disease [108–110]. Our work reveals a new role for TGF-β signals in regulating decision-making, when sensory cues of opposing valance are simultaneously present.

### Specific interneurons modulate 2-nonanone-dependent food leaving in *C. elegans*

The ability to integrate multiple types of sensory stimuli requires not only the responses across peripheral sensory areas, but also the signal processing in downstream network of interneurons [1, 3, 5–8, 111]. In *C. elegans*, a number of sensorimotor responses are modulated by specific contexts via the functions of several interneurons [34, 97, 100, 112]. However, how interneurons mediate decision-making during multisensory behavior is not fully characterized. Here, by examining a number of interneurons that are downstream of the sensory neurons that detect the food-related cues and the repulsive odorant, we find that the AIY interneuron and command interneurons, as well as motor neurons, play a modulatory role in 2-nonanone-dependent food leaving. Disrupting the function of AIY significantly delays the decision to leave the food paired with 2-nonanone, without altering the response to either of the two cues that is presented alone. The AIY interneuron receives synaptic inputs from sensory neurons that detect olfactory, gustatory and thermal information. Previous studies implicate AIY in integrating simultaneously present aversive and attractive cues in olfactory plasticity and in food and serotonin-dependent modulation of sensorimotor responses [34, 97, 112–114]. We propose that AIY may act as an integrating site that receives and processes signals from the food and the repellent 2-nonanone during multisensory integration. In contrast, we did not detect a role for the interneuron AIB with our assay, suggesting the functional diversity among the interneurons in modulating 2-nonanone-dependent food leaving. Our study also implicates the *glr*-*1*- and *nmr*-*1*-expressing neurons in promoting the repellent-dependent food leaving. It is conceivable that some of the *nmr*-*1*-expressing command interneurons and the glr-1-expressing command interneurons or head motor neurons may serve as the downstream-modulated targets for the integrated behavioral response.

### A common set of modulators regulate repellent-dependent food leaving

For freely feeding animals, such as *C. elegans*, appropriate behavioral responses to food sources paired with other sensory cues are critical for survival, because food can be easily contaminated with toxins. To understand to what extent the identified modulators generally regulate integrated responses to foods and repellents, we paired the *E. coli* strain OP50 with either 100% 2-nonanone or 100% benzaldehyde. We also paired 100% 2-nonanone with a second food, *Comamonas.* Interestingly, we found that the TGF-β/DAF-7 pathway and the ASK sensory neuron regulate the integrated responses to these two different pairs of foods and repellents. Avoidance of both 100% 2-nonanone and 100% benzaldehyde depends on the function of the olfactory sensory neurons AWB [43, 45, 91, 115]. Meanwhile, *Comamonas sp* also serves as an attractive food source to the worms [41]. It is conceivable that a common set of modulators represent the contexts where the worm needs to evaluate the opposing values provided by a source of nutrients and a potential threat to generate a behavioral decision.

## Methods

### Strains

*C. elegans* strains were cultivated under the standard conditions [116]. Hermaphrodites were used in this study. The strains that were used in the study include: PR680 *che*-*1(p680)I*, CX14394 *npr*-*5(ok1583)V*, MT15434 *tph*-*1(mg280)II*, CB1112 *cat*-*2(e1112)II*, *MT9455 tbh*-*1(n3247)X*, *RB1161 tbh*-*1(ok1196)X*, *RB993 tdc*-*1(ok914)II*, *MT13113 tdc*-*1(n3419)II*, DR40 *daf*-*1(m40)IV*, PR691 *tax*-*2(p691)I*, *PR671 tax*-*2(p671)I*, RB859 *daf*-*22(ok693)II*, *OH8 ttx*-*3(mg158)X*, *MT150 egl*-*3(n150)V*, CX4 *odr*-*7(ky4)X*, *CX03572 nlp*-*9(tm3579)V*, ZC2685 *npr*-*2(ok419)IV*, VC48 *kpc*-*1(gk8)I*, RB1341 *nlp*-*1(ok1470)X*, RB1289 *npr*-*18(ok1388)X*, CB1372 *daf*-*7(e1372)III*, *ZC2673 gcy*-*33(ok232)V*, *SM2322 daf*-*7(ok3125)III*, AX1295 *gcy*-*35(ok769)I*, QZ81 *ins*-*6(tm2416)II*, QZ126 *ins*-*7(tm2001)IV*, FX02105 *nlp*-*24(tm2105)V*, *RB1902 flp*-*19(ok2460)x*, CX10 *osm*-*9(ky10)IV*, FX02984 *nlp*-*7(tm2984)X*, RB1161 *tbh*-*1(ok1196)X*, RB993 *tdc*-*1(ok914)II*, *KQ361 tdc*-*1(ok914)II*; *daf*-*7(e1372)III*, *KQ363 tdc*-*1(ok914)II*; *daf*-*1(m40)IV*, *KQ364 daf*-*1(m40)IV*; *tbh*-*1(ok1196)X*, *KQ362 daf*-*7(e1372)III*; *tbh*-*1(ok1196)X*, *ZC1952 yxIs25[Pttx*-*3::TeTx::mCherry; Punc*-*122::gfp]*, KQ280 *daf*-*1(m40)IV*; *ftEx98[Pdaf*-*1::daf*-*1::gfp*; *Podr*-*1::dsRed]*, KQ380 *daf*-*1(m40)IV*; *ftEx205*[*Ptdc*-*1::daf*-*1:.gfp*; *Podr*-*1::dsRed*], KQ252 *daf*-*1(m40)IV*; *ftEx70*[*Pbbs*-*1::daf*-*1::gfp*; *Podr*-*1::dsRed*], ZD736 *daf*-*7(ok3125)III*;*qdEx44[Pstr*-*3p::daf*-*7*; *Pges*-*1::gfp]*, ZD729 *daf*-*7(ok3125)III*;*qdEx37[Pdaf*-*7::daf*-*7; Pges*-*1::gfp]*, PY7502 *yxIs34[Pceh*-*36∇::TU813; Pceh*-*36∇:.TU814; Psrtx*-*1::gfp; Punc*-*122::dsRed]*, ZC2393 *yxEx1248 [Pttx*-*3::twk*-*18(gf)::mCherry; Punc*-*122::RFP]*, CX14848 *kyEx4866[Pinx*-*1 ::HisCl1::SL2mCherry; Punc*-*122::dsRed*], CX16040 *kyEx5464[Ptdc*-*1::HisCl1::SL2mCherry]*, ZC1451 *yxEx699[Pnmr*-*1::TeTx::mCherry; Punc*-*122::dsRED]*; QS4 *qrIs2[Psra*-*9::mCasp1; Psra*-*9::gfp; Pelt*-*2::gfp]*, PS6025 *qrIs2[Psra*-*9::mCasp1; Psra*-*9::gfp; Pelt*-*2::gfp];* ZC1552 *yxEx749[Pglr*-*1::TeTx::mCherry; Punc*-*122::gfp]*, PY7505 *oyls84[Pgpa*-*4::TU813; Pgcy*-*27::TU814; Pgcy*-*27::gfp; Punc*-*122::dsRed]*, CX3830 *kyIs102V; kyIs104[Pstr*-*1::mec*-*4(d); Pstr*-*1::gfp];* CX14637 *kyEx4779[Pinx*-*1::TeTx::mCherry; Punc*-*122::gfp]*, CX14993 *kyEx4962[Ptdc*-*1 ::TeTx::mCherry]*, AX2051 *Ex[Pgcy*-*33::egl*-*1; Punc*-*122::dsRed]*, *CX12330 Ex[Psre*-*1 ::TeTx::mCherry; Punc*-*122:RFP]*, *CX7102 lin*-*15B(n765)X; qaIs2241[Pgcy*-*36::egl*-*1; Pgcy*-*35::gfp; lin*-*15(*+)], *ZC2752 nlp*-*7(tm2984)X*; *yxEx1420[Pnlp*-*7::nlp*-*7; Punc*-*122::gfp]*, *ZC2731 daf*-*7(e1372)III; yxEx1409[Pcat*-*2::daf*-*7; Punc*-*122::gfp]*; *ZC2734 daf*-*7(e1372)III*, *yxEx1412[Pocr*-*4::daf*-*7; Punc*-*122::gfp]*

### Behavioral assay for multisensory integration

On a 5 cm-diameter NGM (Nematode Growth Medium) plate, 20-25 young adult worms were placed on a small 1 cm-diameter round-shaped bacterial lawn made of freshly cultivated *E. coli* OP50 strain and left to acclimatize on the lawn for 1-2 hour. Next, a drop of 1 μl 2-nonanone (Sigma Aldrich, Cat # 821-55-6), either 10% (v/v in 100% ethanol) or 100%, was placed on the right-hand side of the lawn and 3 mm away from the lawn. The number of worms on the lawn was counted every 5 minutes for a total of 60 minutes, and the percentage of worms outside the lawn was calculated (Figure 1A and 1B). In some assays, 1 μl of 100% benzaldehyde (Sigma Aldrich, Cat # 100-52-7) was used, instead of 2-nonanone. The OP50 culture was prepared freshly each day by culturing at 27°C for 12-15 hours in NGM medium. For assays using *Comamonas sp* for the food lawn, the experiments were performed in the same way, except that the bacterial strain was cultured with Luria Broth. To determine the time taken to reach the edge of the food lawn, the food lawn was divided into 5 columns with each being 2 mm wide (S5 Fig). The time taken for 90% of the worms to crawl into the column furthest away from the repellent was recorded. Mutants were compared with wild-type animals tested in parallel, and transgenic animals were compared with non-transgenic siblings or wild-type animals tested in parallel on the same day.

The bar graphs in the figures report the percentage of worms outside the lawn at the time point when the significant difference between the tested genotypes was first observed. When there was no significant difference, the bar graphs report the percentage of worms outside the lawn 15 minutes after the start of the assay.

### Transgenes and transgenic animals

To generate a *nlp*-*7* genomic rescue fragment, a 4.7 kb PCR product was amplified from genomic DNA that included 2.5 kb 5’ upstream sequence, the *nlp*-*7* coding region, and 1 kb 3’ downstream sequence (NLP-7F: 5’-CATGTTTTTGATCATTTTCGAAC-‘3 and NLP-7R3’UTR: 5’-AATATCGTATGCCAACTTGAAC-‘3). The *nlp*-*7* genomic PCR product was injected into the *nlp*-*7(tm2984)* animals. To generate the construct expressing a wild-type *daf*-*7* cDNA in the OLQ or ADE sensory neurons, the *daf*-*7* cDNA was amplified from PJM016 (Gift from Dr. Dennis Kim and Dr. Joshua Meisel [71]). The *daf*-*7* cDNA product was cloned into a gateway destination vector that contained an *unc*-*54* 3’UTR using the Nhe-1 and Kpn-1 sites. Both the promoter regions of *ocr*-*4* (4.0 kb promoter for expression in OLQ) and *cat*-*2* (1.1 kb promoter for expression in ADE) were amplified from genomic DNA (CAT-2F: CTAGCAGGCCCAATCTTTTCTG and CAT-2R: TCCTCTTCCAATTTTTCAAGGGGT/OCR-4F: 5’-TTCTAATATTGCTCCATCAAC-‘3 and OCR-4R: 5’-TAATACAAGTTAGATTCAGAGAATA-‘3) and cloned into the entry-TOPO vector PCR8 (Invitrogen). The expression clones, *Pcat*-*2::daf*-*7* and *Pocr*-*4::daf*-*7*, were generated using LR recombination reactions (Invitrogen). Each transgene was injected at 30-50 ng/μl with the co-injection marker as previously described [117].

### Lawn-leaving assay

Lawn-leaving assay was performed and analyzed similarly as the assay for multisensory integration, except that no repulsive chemical was present. Briefly, animals were placed on a 1 cm-diameter round-shaped bacterial lawn of OP50 and left for 10 minutes to acclimatize before examining food leaving over a period of one hour by counting the number of worms that were present on the food lawn every 5 minutes for a total of 60 minutes.

### 2-nonanone avoidance assay

To examine the avoidance of 2-nonanone, chemotaxis assays were performed essentially as previously described [43]. Briefly, animals were placed in the center of a square plate that was divided into sectors A - F and 2 drops of 1 μl of 2-nonanone was added to one side and 2 drops of 1 μl ethanol was added to the opposite side of the plate as control. Approximately 100 worms were used in each assay. Chemotactic avoidance was analyzed by counting the number of worms in the sectors A-B, C-D and E-F with E-F being furthest away from the 2-nonanone point sources (S5 Fig). The avoidance index was calculated as the number of animals in sectors A and B minus the number of animals in the sectors E and F and normalized with the total number of animals in all 6 sectors on plate.

## Acknowledgements

We would like to thank Drs. Kevah Ashrafi, Cori Bargmann, Dennis Kim, Richard Komuniecki and Piali Sengupta for sharing reagents. We also thank members of the Zhang lab for insightful discussion for this project. The research in the Zhang lab is supported by Harvard University (PRISE program for G.L.) and The National Institutes of Health (NIH).

**The authors declare no competing interest.**

## Supporting information

**S1 Table. 2-nonanone avoidance assay.**

Wild type, mutants and transgenic animals are examined for avoiding 100% 2-nonanone as previously described (Troemel et al., 1997 and S5 Figure). Avoidance Index was calculated as described in S5 Figure. The avoidance in each genotype is represented by the average avoidance index of individual assays. n = 2 – 4 assays each genotype, 75-100 animals tested in each assay, mutants and transgenic animals are compared with wild-type animals or the non-transgenic siblings tested on the same days with student’s *t* test, Mean ± SEM.

**S2 Table. Time to reach the edge of the food lawn during multisensory integration**

Wild type, mutants and transgenic animals are examined for the time taken to reach the edge of the food lawn away from the repellent. The average time taken for 90% of the worms in one assay to reach the edge of a *E. coli* OP50 food lawns during exposure to 100% 2-nonanone is presented for each genotype (Experimental Procedures). n = 2 – 4 assays for each genotype, 20-25 animals tested in each assay, mutants or transgenic animals are compared with wild-type animals or non-transgenic siblings tested on the same days with student’s *t* test, Mean ± SEM.

**S3 Table. Spontaneous food leaving from an OP50 lawn without pairing with 2-nonanone.**

Wild type, mutant animals and transgenic animals are examined for food leaving for 1 hour. Young adult worms are placed on an OP50 food lawn for 1 hour and the number of worms on food lawn is counted every 5 minutes for a total assay time of 60 minutes. The percentage of worms off the food lawn at 15 minutes is reported. n = 2 – 4 assays for each genotype and 20-25 worms in each assay, mutants or transgenic animals are compared with wild-type animals tested in parallel with student’s *t*-test, Mean ± SEM.

**S4 Table. Many signaling mutants show no phenotype in 2-nonanone-dependent food leaving.** Wild type, mutant and transgenic animals are examined for leaving an *E. coli* OP50 food lawn paired with 100% 2-nonanone. The average percentage of worms outside the food lawn at 15 minutes is reported. Mutants or transgenic animals are compared with the wild-type control tested on the same days with student’s *t* test, n = 2 – 4 assays for each genotype, 20-25 animals in each assay, Mean ± SEM.

**S5 Fig. Schematics of assays**

A. Assay to measure time taken to reach the edge of the food lawn (Experimental Procedures).
B. Chemotaxis assay for avoidance of 100% 2-nonanone (Experimental Procedures)

**S6 Fig. Additional alleles of *tdc*-*1* and *tbh*-*1* mutants are also wild-type for 2-nonanone-dependent food leaving.**

Each bar graph shows the percentage of animals outside the food lawn 15 minutes after the start of the assay, mutants are compared with wild type tested in parallel with Student’s *t* test, n= 3 assays each; mean ± SEM, n.s., not significant.

**S7 Fig. DAF-7 produced by the sensory neurons ASI does not regulate 2-nonanone-dependent food leaving.**

Expressing a wild-type *daf*-*7* cDNA in the sensory neurons ASI does not rescue the delayed decision phenotype in the *daf*-*7(e1372)* mutant animals (n = 4, 3 and 4 assays for wild type, transgenic animals and non-transgenic siblings, respectively).

Each bar graph shows the average percentage of worms outside the lawn 15 minutes after the start of the assay, transgenic animals are compared with non-transgenic siblings using Student’s *t-test*, n.s., not significant.

**S8 Fig. Expressing the** *daf*-*1* **cDNA in sensory neurons with** *osm*-*6* **promoter does not rescue the delayed-decision phenotype in the** *daf*-*1(m40)* **mutants.** The transgenic animals (n = 3 assays) are compared with non-transgenic siblings (n = 4 assays) with Student’s *t* test, wild type = 3 assays; bar graph shows the percentage of worms outside of lawn 15 minutes after the start of the assay, mean ± SEM, n.s., not significant.

**S9 Movie. Wild-type worms performing food leaving on an** *E. coli* **OP50 lawn paired with 100% 2-nonanone.**

**S10 Movie. Wild-type worms performing food leaving on an** *E. coli* **OP50 food lawn paired with 100% benzaldehyde.**

